# Candidate regulators and target genes of drought stress in needles and roots of Norway spruce

**DOI:** 10.1101/517151

**Authors:** Julia C. Haas, Alexander Vergara, Vaughan Hurry, Nathaniel R. Street

## Abstract

Drought stress impacts on seedling establishment, survival and whole-plant productivity. Drought stress responses have been extensively studied at the physiological and molecular level in angiosperms, particularly in agricultural species and the model *Arabidopsis thaliana*, with the vast majority of work performed on aboveground tissues. Boreal forests are dominated by coniferous tree species and cover vast areas of the terrestrial surface. These areas are predicted to be particularly influenced by ongoing climate change and will be exposed to more frequent and acute drought. The associated impact at all stages of the forest tree life cycle is expected to have large-scale ecological and economic impacts. To provide a comprehensive understanding of the drought response mechanisms of *Picea abies* seedlings, we assayed the physiological response of needles and transcriptional responses of roots and needles after exposure to mild and severe drought. Shoots and needles showed extensive reversible plasticity for physiological measures indicative of drought response mechanisms, including stomatal conductance (*g_s_*) and shoot water potential. Root and needle transcriptional responses contrasted, with an extensive root-specific down-regulation of growth. When we compared the responses of *P. abies* with previously-characterised *A. thaliana* drought response genes, we found that the majority of the genes were conserved across lineages. However, in *P. abies*, transcription factors (TFs) previously identified as belonging to the ABA-dependent pathway had a more limited role and most differentially expressed genes were specific to the stress response of *P. abies*. These results highlight the importance of profiling both above- and below-ground tissues and provide a comprehensive framework to advance understanding of the drought response mechanism of *P. abies*.

**One sentence summary:** Analysis of the drought transcriptome of Norway spruce reveals divergent molecular response pathways in conifers.

## Introduction

Boreal forests are dominated by coniferous trees (Shorohova et al., 2011) and represent approximately one third of the terrestrial forest area (FAO, 2010). They act as sinks of atmospheric CO2 and carbon emissions from anthropogenic sources and are important in balancing the global carbon cycle (Kirschbaum and Fischlin, 1996; Pan et al., 2011). Growth conditions in the boreal zone are often limiting for plants, with a cold climate and long periods of sub-zero temperatures leading to short growing seasons (Troeng and Linder, 1982; Burton et al., 2010; Kneeshaw et al., 2011). Recently, climate change has caused a mean annual temperature increase of 1.5 °C in high latitude boreal forests, with temperatures predicted to rise by an additional 5 °C by 2100 (IPCC, 2013). Observations made in boreal forest biomes over Fennoscandia, North America and Russia have identified accelerated growth in response to higher temperatures and longer growing seasons (Lapenis et al., 2005; Zhang et al., 2008; Kauppi et al., 2014). However, such positive growth responses are dependent on adequate water availability and the same studies also reported severe increases in summer drought and evapotranspiration in boreal regions (Zhang et al., 2008; Allen et al., 2010; Beck et al., 2011; Peng et al., 2011). Drought stress is a major cause of tree mortality and a threat to key tree species in boreal forest stands, including Norway spruce (*Picea abies* (L.) H. Karst) in southern Scandinavia (Kellomäki et al., 2008; Ge et al., 2011). To understand how future tree populations may cope with drought, and whether boreal forests will continue to serve as carbon sinks in a future changing climate, studies of phenotypic plasticity for drought stress responses, including transcriptomic changes that facilitate identification of the underlying drought response mechanisms of keystone species such as Norway spruce, are needed.

Drought has always been a constraint for productivity of agricultural crops worldwide and major insights to the biochemical and molecular mechanisms important for plant drought response have been gained from studies of herbaceous plants such as Arabidopsis (*Arabidopsis thaliana* L.), rice (*oryza sativa* L.) and maize (Zea *maize* L.). To avoid or tolerate water deficit, maintenance of cellular turgor is of high priority and plants therefore accumulate metabolites such as sugars, amino acids and amines (Tabaeizadeh, 1998). In addition, cellular components such as proteins and membranes require stabilisation, which involves late embryogenesis abundant (LEA) proteins, heat shock proteins (HSP) and dehydrins, together with the above metabolites, which also function as antioxidants (Harfouche et al., 2014). The phytohormone ABA (abscisic acid) is an important signal leading to the expression of various stress responsive genes and transcriptional networks (Yamaguchi-Shinozaki and Shinozaki, 2006). The ABA signal is perceived by the PYR/PYL/RCAR (pyrabactin resistance/pyrabactin resistance1-like/regulatory component of ABA receptors) receptors and AREB (abscisic acid-responsive element binding proteins)/ABF (ABRE binding factor) TF (transcription factors) induce further gene expression changes after phosphorylation by subclass III SnRK2 (SNF1 related protein kinase) kinases (Cutler et al., 2010; Umezawa et al., 2010). TFs from the MYB/MYC, NAC, WRKY and NF-Y families have additionally been shown to regulate ABA-responsive gene expression under drought stress (Singh and Laxmi, 2015). In addition, an ABA-independent pathway for signal transduction under drought stress exists, involving the DREB (dehydration-responsive element binding protein)/CBF (C-repeat binding factor) TFs, a subfamily of AP2 (APETALA 2)/ERF (ethylene-responsive element-binding factor) transcriptional activators (Nakashima et al., 2009). Due to their role as master switches of gene expression, modifying the expression of TFs has a high capacity to improve drought responses (Rabara et al., 2014). As an example, in transgenic rice, overexpression of *SNAC1* (Stress-Responsive NAC1) was shown to increase drought tolerance and also increased yield in field trials (Hu et al., 2006). Importantly, regulation and signalling of abiotic stress responses in cereals was found to be highly conserved, as demonstrated by improved drought tolerance in wheat lines over-expressing the rice *SNAC1* (Saad et al., 2013). The application of high-throughput RNA-Seq (RNA-Sequencing) to assay drought responses provides a means to further disentangle complex regulatory networks in higher plants and to identify candidate genes for the development of new varieties with improved drought tolerance through targeted genetic transformation, genome editing or breeding.

Physiological changes, such as stomatal closure are the primary cause of drought-induced decreases in productivity. Stomatal closure is stimulated by increasing levels of endogenous ABA during drought stress (Iuchi et al., 2001), which consequently, results in reduced availability of CO2 for photosynthesis. Plants differ in their sensitivity to dehydration with substantial intra- and inter-specific variation in response to drought. Herbaceous and woody angiosperm species are thought to be mainly anisohydric in that they display a risk-taking behaviour, maintaining stomatal conductance (*g_s_*) under drought stress in order to keep productivity at a high level (Sade et al., 2012). In contrast, isohydric species, including most gymnosperms of the *pinophyta* such as Norway spruce, initiate stomatal closure early during drought and also maintain high levels of ABA in the foliage, restricting fast recovery after periods of water deficit (Brodribb and McAdam, 2013; Brodribb et al., 2014). Maintenance of an efficient hydraulic system to ensure transport of water from the soil (xylem) and to reduce water loss from leaves (via stomata) is especially important in in trees, which are long-lived sessile organisms, for optimal plant hydration under seasonally and annually fluctuating environmental conditions (Raven, 1977; Edwards et al., 1998). Differences in the regulation of water status can determine survival of trees in response to drought with improper response mechanisms resulting in mortality from either carbon starvation or hydraulic failure in isohydric and anisohydric species, respectively. It is important to understand how species will respond within this continuum of response mechanisms in light of predicted patterns of intensity, duration and frequency of future drought events (McDowell et al., 2008). This will be increasingly important to forest production as climate change increases the number and duration of drought events, including changes to VPD (vapour pressure deficit), which will particularly reduce productivity in ecosystems where the isohydric *pinophyta* dominate (Roman et al., 2015).

The role of ABA production in in roots and transport to the foliage to actively reduce guard cell turgor is generally accepted as the primary mechanism mediating stomatal closure during drought stress in herbaceous plants (Schroeder et al., 2001; Roelfsema and Hedrich, 2005). However, in trees the time required for transport of ABA to the leaves via the vascular tissue exceeds the observed time needed to close stomata (Zimmermann and Brown, 1971; Schulze and Hall, 1982). The importance of root-derived ABA is therefore questionable (Wilkinson and Davies, 2002) and with the relative importance of the ABA pools in different plant tissues remains unresolved. Nevertheless, ABA has a role in drought signalling in angiosperm tree species, for example in *Populus* species transcriptome studies have identified potential orthologs to Arabidopsis PP2Cs (Protein Phosphatase 2C), functioning in ABA signalling (Chen et al., 2015), as well as putative AREB/ABF homologs, which are responsive to exogenously applied ABA (Ji et al., 2013). In addition, representatives of the ABA-independent pathway, e.g. members of the *DREB2* genes, have been proposed to have similar functions in drought response in poplar species to herbaceous plants (Chen et al., 2011). However, there are examples of considerable variation in sensitivity to ABA and differences in the regulation of gene expression between species even within the same genus (Street et al., 2006). Furthermore, the isohydric stomatal response of gymnosperm species also includes a high sensitivity to ABA (Brodribb and McAdam, 2013; Brodribb et al., 2014) but differences to anisohydric plants in utilising hormonal signals and other molecular mechanisms as effectors of stomatal closure may exist. The molecular basis of isohydrism has been less studied and genes identified based on sequence homology in gymnosperms therefore require functional confirmation in drought stress.

Drought tolerance not only depends on aboveground stomatal behaviour, as roots have an essential role in determining water uptake from the environment and are the first to sense a soil water deficit. Norway spruce is often found to have a shallow root system (Kalliokoski 2011) with the majority of fine roots located in the upper soil layers (Borja et al. 2008, in cermak). As such, Norway spruce, and especially seedlings, are highly sensitive to summers with lower precipitation and high evapotranspiration, which will become more prevalent in as a result of climate change. While large, mature trees with extensive root systems and large stored water reserves can partially buffer the effects of mild drought, seedlings lack this capacity (Williams, 1997; Sparks and Black, 1999). Despite the essential role of roots in determining the physiological outcome of drought stress, research has mainly focused on aboveground responses and regulatory mechanisms. Roots may, however, induce specific regulatory stress response mechanisms in response to early perception of soil drying, even in response to moderate drought. This is supported by root-specific up-regulation of drought response genes including *DREB1*, another representative of the ABA-independent pathway, in roots of loblolly pine (Lorenz et al., 2011), poplar (Cohen et al., 2010) and soybean (Ha et al., 2015) and higher up-regulation in maize roots than leaves (Liu et al., 2013).

In this study, RNA-Sequencing was used to assay transcriptional changes induced under controlled, gradual soil water depletion and after recovery in three-year old Norway spruce seedlings. Needles and roots of seedlings experiencing a mild drought stress at FC (field capacity) of 60, 40 and 30% and after a more extreme water stress that resulted in stomatal closure were compared to well-watered (80% FC) samples, in addition to samples after re-irrigation. There was extensive reversible plasticity for physiological measures indicative of drought response mechanisms, including *gs* and shoot water potential. Needles and roots showed contrasting transcriptome remodulation in response to drought, including differences in plasticity upon rewatering. Additionally, the conservation of drought response regulation between Norway spruce, as a representative conifer, and Arabidopsis, as a representative angiosperm, was explored. Although there was a set of previously characterised drought responsive genes identified in angiosperms that have conserved drought response in Norway spruce, there was also extensive divergence in transcriptional response between the lineages. These results advance understanding of the transcriptional contribution to drought response in conifers and highlight the contrasting transcriptional responses of roots and needles.

## Results

### The physiological drought response of Picea abies seedlings

An experiment was conducted using three-year-old seedlings of Norway spruce to characterise physiological and transcriptional responses after *exposure* to a short-term mild and severe drought and subsequent recovery after re-irrigation. Physiological stress of seedlings from well-watered, mild drought stress (days 2, 4 and 5), severe drought stress (days 18 and 21) and recovery (four days after re-irrigation) conditions was determined by measuring midday shoot water potential. The same seedlings were sampled for transcriptome profiling. In addition, stomatal conductance (gs) and photosynthetic activity (Amax) of a set of well-watered and treated plants was measured over the course of the experiment. Treatment severity increased during the first five days, with a 10% decrease in field capacity (FC) each day until 30% (Fig. 1A), after which FC was maintained at 30% for seven days. Thereafter, the seedlings were exposed to a more severe drought stress by withholding water until catastrophic dysfunction of g_s_ and A_max_ was detected in aboveground plant tissues (Fig 1B & 1C). g_s_ decreased within the first five days, remained stable at 30% FC and significantly decreased in response to imposition of the severe drought stress (p < 0.01), when soil water content was 20% FC (Fig. 1B). Photosynthetic CO_2_ assimilation (A_max_) was unaffected until imposition of the severe treatment, after which there was an abrupt decline at day 17 (p < 0.01) in response to closure of stomata (Fig. 1C). Shoot water potential changed significantly after five days of mild water stress (p < 0.05) (Fig. 1D), but temporarily recovered during the phase when FC was held at 30%. After initiation of the severe drought stress, shoot water potential showed a further significant decrease (p < 0.01). These results demonstrated that Norway spruce is an isohydric conifer species, reducing gs over a very narrow range of water potential (between −1.1 and −1.8 MPa) in order to maintain midday shoot water potential at a reduced but relatively constant level (−2.1 MPa). Both g_s_ and A_max_ were maximally limited after 18 days, five days after initiation of the severe drought stress. Seedlings in the severe drought treatment could be recovered by re-watering after day 21 (Fig. 1D). However, continuation of the severe drought for an additional two days resulted in defoliation and death of the seedlings (not shown). Four days after re-watering (at day 21), shoot water potential returned to pre-stress levels, g_s_ recovered to ~60% and A_max_ to ~80% of the levels prior to initiation of the drought treatment. g_s_ and A_max_ of control seedlings was unchanged throughout the period of the experiment (Fig. 1B & 1C).

**Figure 1.**
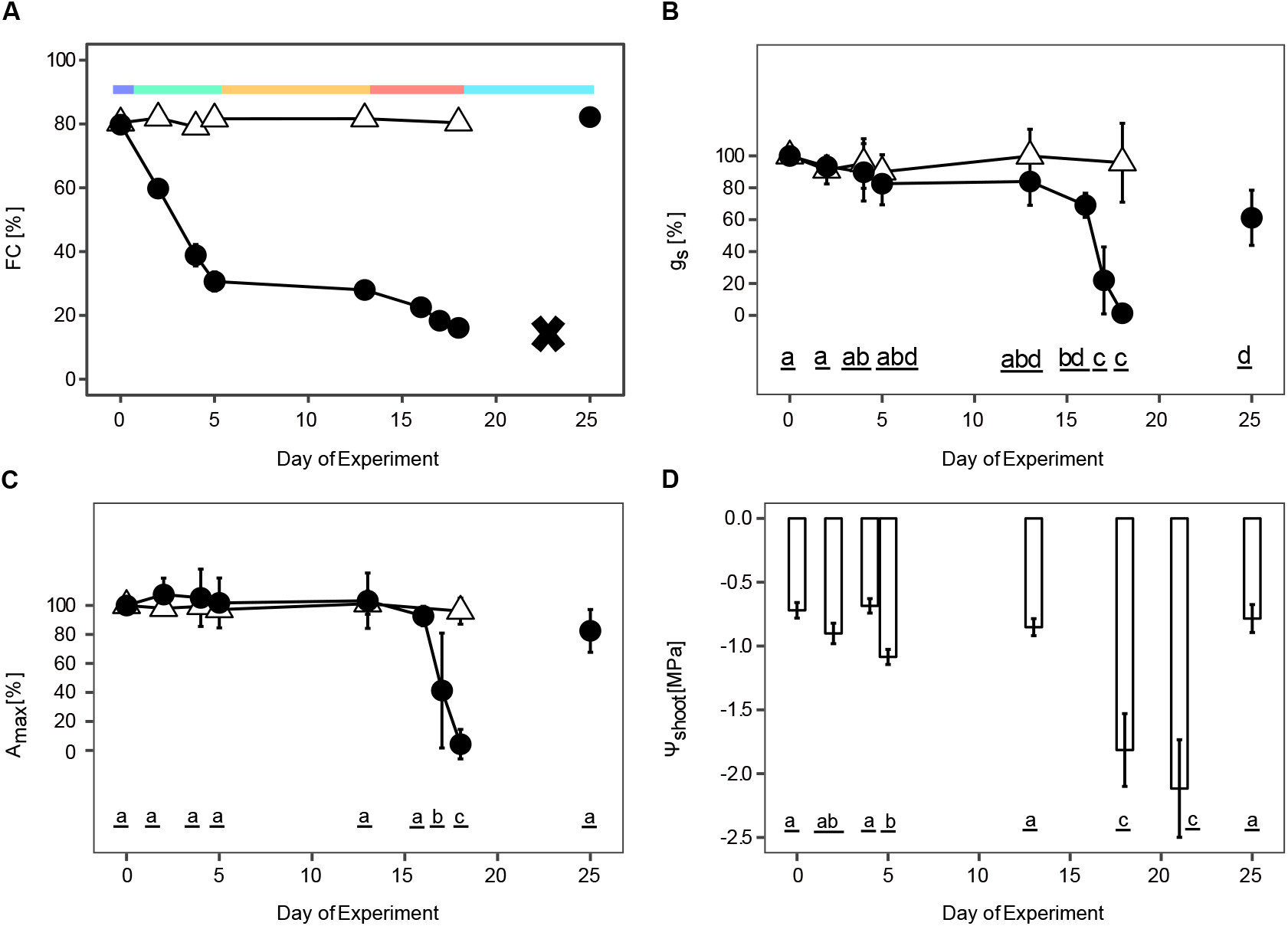
Physiological response of Norway spruce seedlings to drought. Norway spruce seedlings were subjected to increasing levels of water deficit (filled circles) or kept in well-watered control conditions (clear triangles). A) Soil moisture dropped from 80 % (blue) to 30% field capacity (FC) after withholding water for five days and was regarded as mild water deficit (green). After maintaining this condition for seven days (orange), a more severe drought stress was induced (red), again by withholding water. After 21 days water stressed plants were rehydrated (turquoise) and soil moisture levels recovered to the level of well-watered plants. B) g_s_ and C) photosynthetic carbon assimilation (A_max_) were measured continuously on three control and drought stressed plants. D) Midday water potential (ψ_shoot_) was measured on shoots of three to five independent plants per time point and statistics performed on the average values of three measurements per plant. Data are means and error bars represent the ± 95% confidence intervals. Letters above bars represent statistically significant differences (p < 0.05) between time points of the drought stress.

### Needles and roots have distinct transcriptional responses to drought

Comparison of drought stressed to control needle and root samples revealed only limited changes in relative transcript abundance in response to the mild drought stress (two, four and five days without water) while a more extensive remodulation of the transcriptome was induced by the severe drought stress, particularly in roots (18 and 21 days after treatment initiation) (Fig. 2A & S1). Re-irrigation reversed the transcriptional changes in needles, but the transcriptome of roots remained distinct after recovery from that of control and drought samples (Fig. 2A & S1B). In total, a greater number of transcripts were DE (differentially expressed) in roots than in needles (6194 in comparison to 1403 DE genes, respectively, of which 636 were commonly DE in both tissues), indicating that roots may have experienced more severe water stress than the needles (Fig. 2B) or that response mechanisms under transcriptional control are more important to the drought response of roots of Norway spruce seedlings. Mild drought induced DE of 98 genes in needles, of which 91 were up- and seven down-regulated (Fig. 2C and S2), and similarly 98 genes were DE in roots, with 90 up- and eight down-regulated genes (Fig. 2C and S2). In contrast, 1304 genes were DE in response to severe drought stress in needles, with 871 of these up- and 433 down-regulated, while in roots 5835 genes were DE in response to severe drought stress, of which 1926 DE genes were up- and 3909 down-regulated. Extensive down-regulation was therefore an important component of the transcriptional response mechanism of roots during severe drought stress and active remodulation of the transcriptome was far more prominent in roots than in needles. After re-irrigation, 23 genes were DE in needles, 14 up- and 9 down-regulated, whereas in roots 853 genes were DE, 603 up- and 250 down-regulated.

**Figure 2.**
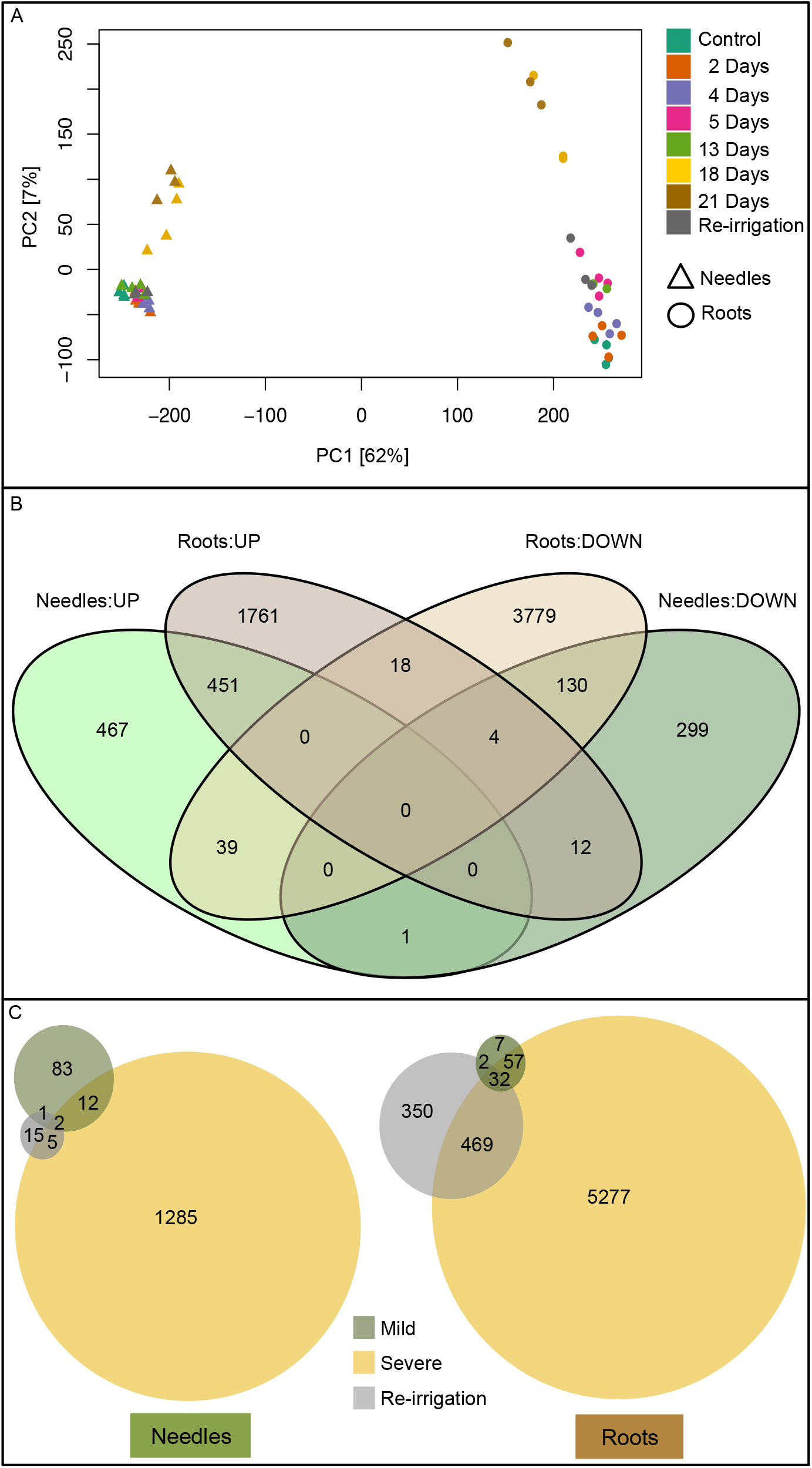
Effect of drought on the transcriptomes of needles and roots in Norway spruce. Principal component analysis (PCA) plot of transcriptomic data of drought stressed needle and root (A) samples. Expression data of control, mild and severe drought stressed seedlings and after rehydration was normalised using variance stabilising transformation before ordination analysis. Three to four plants were sampled at each time point and used for RNA sequencing. The first two components of the PCA are shown with samples colored by soil water percentage field capacity. B) The total number of genes up- and down-regulated in needle (light green, up-regulated in needles; dark green, down-regulated in needles) and root (light brown, up-regulated in roots; dark brown, down-regulated in roots) samples that were differentially expressed. C) Venn-diagrams of differentially expressed genes in needles (left) and roots (right), comparing the mild (yellow) and severe (dark green) drought stage and after rehydration (gray) separately against well-watered expression levels.

### The transcriptional response of roots involves extensive down-regulation of growth-related processes

To explore the biological processes involved in the transcriptome response to drought, 13 clusters representing co-expressed sets of genes were identified and functional annotations assigned to these (Fig. 3A & 4A and Supplementary file 1 & 2 for gene clusters primarily up- or down-regulated in response to drought, respectively). Gene Ontology (GO) terms could be assigned to ~34% of the DE genes and GOSlim terms were used to provide a global overview of the biological processes active in response to drought in needles and roots (Fig. 3B and 4B). The GOSlim term “response to stress” (GO:0006950) was significantly enriched (p < 0.05) in cluster 3 (Fig. 3B), which contained genes commonly up-regulated in both needles and roots in response to severe drought. Cluster 10 (Fig. 4A) comprised a large set of genes that were specifically down-regulated in roots during severe drought, with this down-regulation then reversed upon re-irrigation, indicating recovery from water stress. This cluster of genes was significantly enriched (p < 0.05) for GOSlim terms associated with growth including “anatomical structure development” (GO:0048856), “transport” (GO:0006810), “reproduction” (GO:0000003), “carbohydrate metabolic process” (GO:0005975), “cell wall organisation or biogenesis” (GO:0071554), “DNA metabolic process” (GO:0006259), “chromosome organisation” (GO:0051276), “cell differentiation” (GO:0030154), “cell cycle” (GO:0007049), “growth” (GO:0040007), “anatomical structure formation involved in morphogenesis” (GO:0048646), “cell morphogenesis” (GO:0000902), “developmental maturation” (GO:0021700), “cell division” (GO:0051301), “cell proliferation” (GO:0008283) and “pigmentation” (GO:0043473). As such, active down-regulation of growth appeared to be an important component of the drought response mechanisms in the roots of Norway spruce seedlings. Finally, “response to stress” (GO:0006950) was significantly enriched (p < 0.05) in cluster 11 (Fig. 4B), combining DE genes down-regulated in roots both during severe drought and after re-irrigation. Cluster 13 (Fig. 4A) contained DE genes up-regulated during mild stress in needles, down-regulated in roots and needles during severe drought and with expression that returned to near-control levels after re-irrigation. These genes were significantly enriched (p < 0.05) for the term “cell wall organisation or biogenesis” (GO:0071554). Taken together, these results indicate that growth processes were actively down-regulated in roots in response to drought with this transcriptional response being highly plastic, showing rapid reversal after re-irrigation.

**Figure 3.**
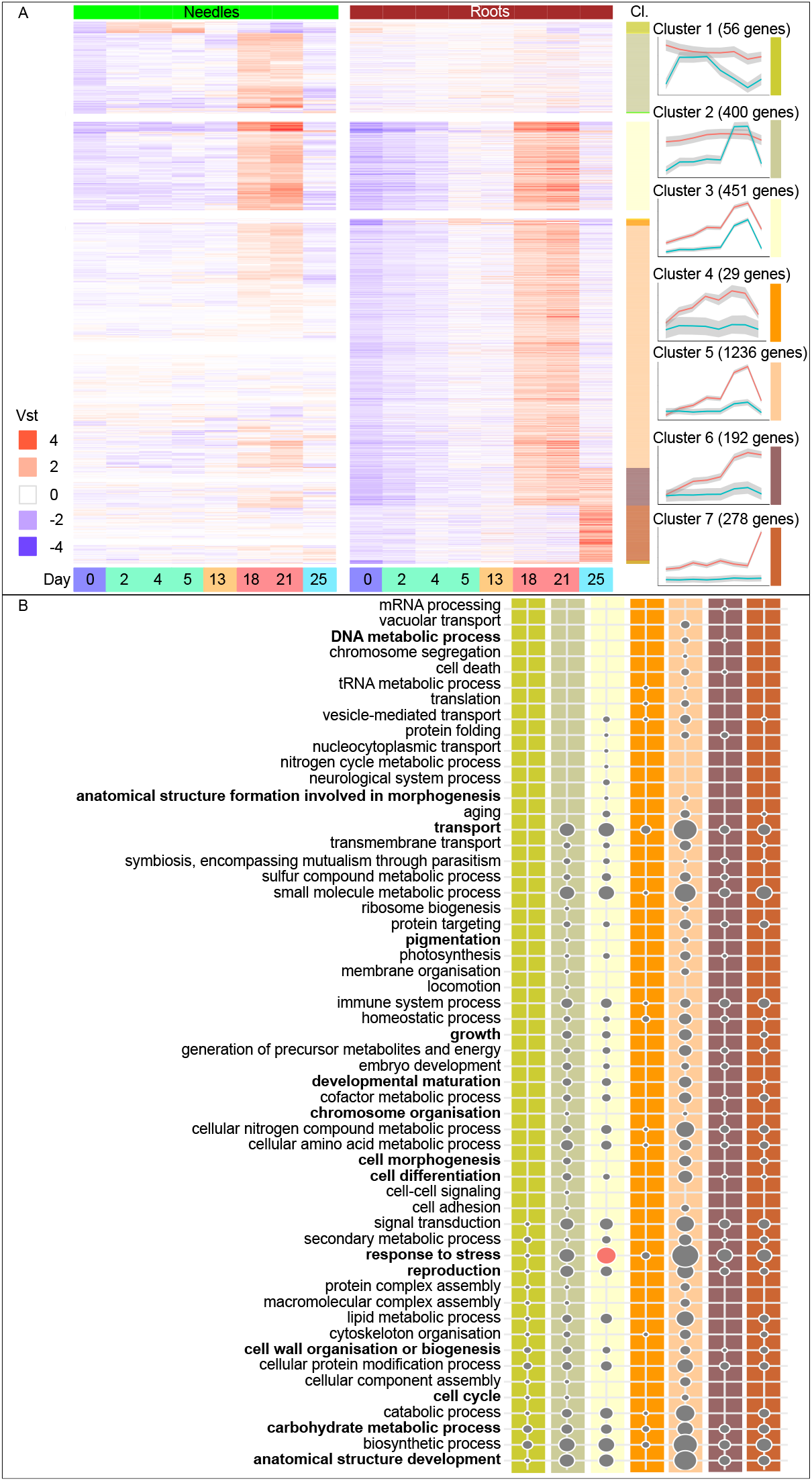
Gene Ontology Slim enrichment analysis of biological processes up-regulated in response to drought stress. Heatmaps of differentially expressed genes with expression over the eight sampling time points (day 0, 2, 4, 5, 13, 18, 21 and 25), separated by tissue, needles (green) and roots (brown). Time points were coloured corresponding to Figure 1 in blue at day 0, green at day 2,4 and 5, orange at day 13, red at day 18 and 21 and turquoise at day 25. A) Up-regulated genes of needle (cluster 1 and 2) and root specific (cluster 4 to 7) clusters or in common between the tissues (cluster 3). Displayed are the vst values scaled by row means. The seven highly populated clusters (Cl.) are detailed on the side and the average trend of gene expression and a 95% confidence interval indicated for both tissues. B) The lower panel of the figure describes all Gene Ontology Slim biological processes categories represented in the up-regulated genes sorted by the heatmap clusters. Circle sizes are relative to the top category; “response to stress” with 76 up-regulated genes. Circles in red color represent significantly enriched categories (False Discovery Rate adjusted p-value < 0.05).

**Figure 4.**
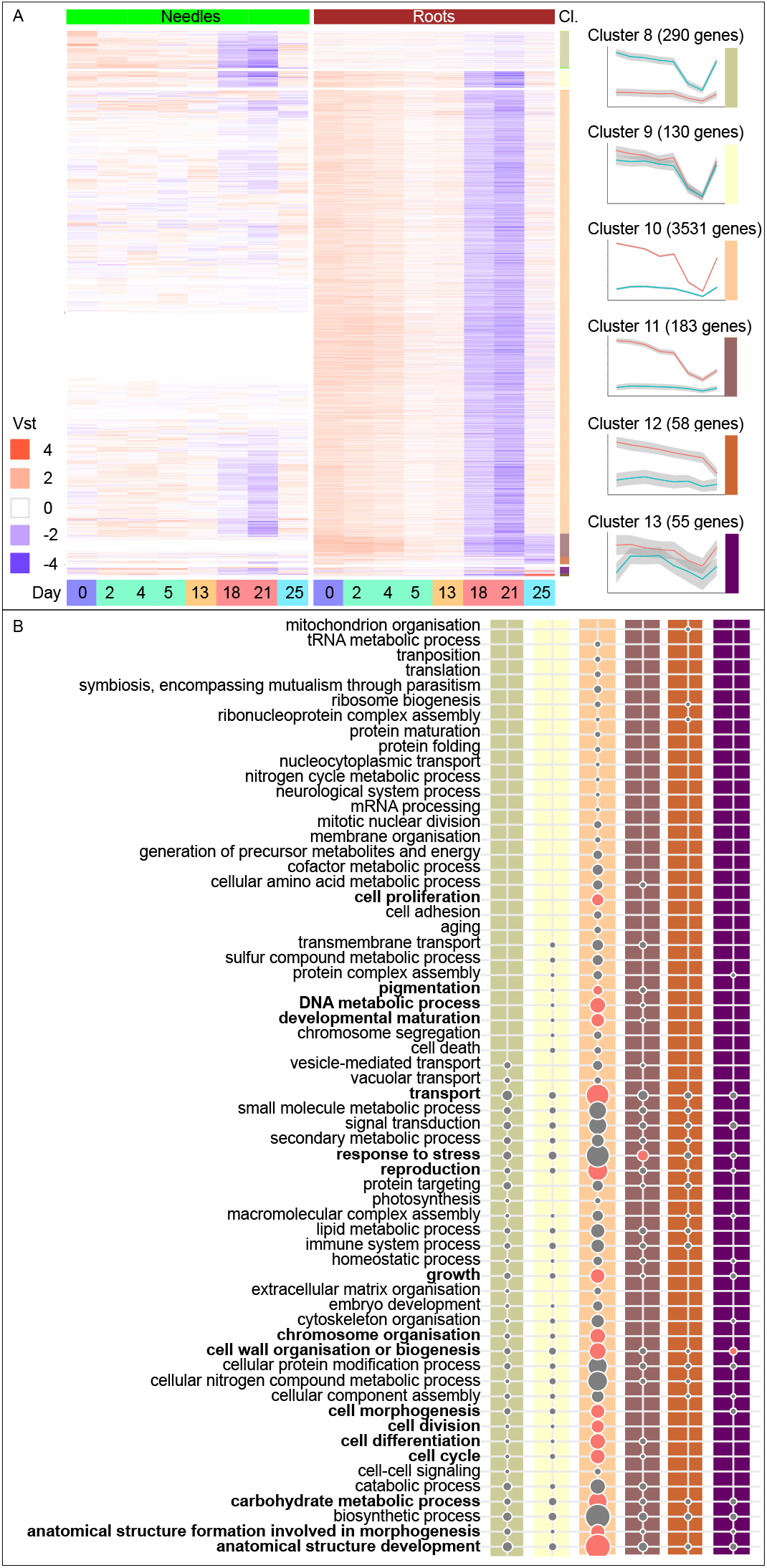
Gene Ontology Slim enrichment analysis of biological processes down-regulated in response to drought stress. Heatmaps of differentially expressed genes with expression over the eight sampling time points (day 0, 2, 4, 5, 13, 18, 21 and 25), separated by tissue, needles (green) and roots (brown). Time points were coloured corresponding to Figure 1 in blue at day 0, green at day 2,4 and 5, orange at day 13, red at day 18 and 21 and turquoise at day 25. A) Down-regulated genes of needle (cluster 8) and root specific clusters (10 to 12) or common between the tissues (cluster 9 and 13). Displayed are the vst values scaled by row means. The six highly populated clusters (Cl.) are detailed on the side and the average trend of gene expression and a 95% confidence interval indicated for both tissues. B) The lower panel of the figure describes all Gene Ontology Slim biological processes categories represented in the down-regulated genes sorted by the heatmap clusters. Circle sizes are relative to the top category; “anatomical structure development” with 220 down-regulated genes. Circles in red color represent significantly enriched categories (False Discovery Rate adjusted p-value < 0.05).

In agreement, re-irrigation resulted in up-regulation (p > 0.05) in roots of metabolism (e.g. “small molecule metabolic process” (GO:0044281), “carbohydrate metabolic process” (GO:0005975) and “biosynthetic process” (GO:0009058) (Fig. 3B, cluster 7)), indicating an ongoing recovery from the severe stress. However, “response to stress” (GO:0006950) remained induced in roots (p > 0.05) after re-irrigation (Fig. 3B, cluster 6). Re-irrigation also induced additional genes associated with a stress response to hypoxia in roots (Fig. 3A, cluster 7) (e.g. *ADH1* and *LOX2*, see Supplementary file 3 for details of genes), suggesting that a high level of responsiveness was maintained in roots even after release of drought stress.

### Functional analysis of the Norway spruce transcriptome in comparison to Arabidopsis

Arabidopsis is the most comprehensively characterised species for a range of abiotic stresses, including drought. To ascertain how relevant biological annotation of genes in this model system is for informing studies in Norway spruce, sequence homology analysis was performed. Arabidopsis orthologs were identified for 13%(184 of 1400) and 14% (888 of 6160) of DE genes in needles and roots, respectively, and homologs identified for 57% and 55%, respectively (Table 1 and Supplementary file 3). An additional 18% and 19%, respectively, were homologous to gymnosperm or other angiosperm species available at the PLAZA resource (Proost et al., 2015) and only 12% (166 in needles and 762 in roots) of the drought responsive genes had no homology match and were termed Norway spruce-specific singletons. There was no significant enrichment for singletons within the DE genes. As such, the observed transcriptional drought response of Norway spruce seedlings primarily involved a set of genes that are broadly-conserved across lineages.

**Table 1.** Percentage of drought responsive transcripts in Norway spruce with sequence conservation in plants.

|  | NeedleDE [%] | RootDE [%] | Common |
| --- | --- | --- | --- |
| Homology (DE) | (1400) | (6160) | (634) |
| Ortholog (Ath) | 13.1 (184) | 14.4 (888) | (82) |
| Homolog (Ath) | 57.1 (800) | 54.7 (3369) | (369) |
| Angiosperm | 0.0 (0) | 0.1 (4) | (0) |
| Gymnosperm | 15.7 (220) | 15.7 (970) | (100) |
| Others | 2.1 (30) | 2.7 (167) | (13) |
| Singletons (Pab) | 11.9 (166) | 12.4 (762) | (70) |

To specifically examine whether extensively characterised Arabidopsis drought-response genes displayed similar transcriptional response to drought in Norway spruce, a manually-curated set of 373 genes was further considered (Supplementary file 3). Orthologs or homologs were identified for 335 (89.9%) of these genes in the Norway spruce genome, represented by 979 Norway spruce gene models (Table 2 and Fig. S3), with > 87 % of these expressed in the transcriptomes of the current control and drought stressed needle and root samples. Only a subset was DE in needles (10.5 %; 39 genes) or roots (31.4 %; 117 genes) and only this subset was therefore considered to have evidence of conserved function (indicated by transcriptional response) in response to drought stress in Norway spruce seedlings (Table 2, Fig. S3 and Supplementary file 3). These DE genes were assigned into the clusters detailed in Fig. 3A and 4A based on their expression profiles (Supplementary file 3). Among this set of genes, conservation existed for genes functioning in induction of solute accumulation, such as for sugars and amino acids, (e.g. *GolS1, RS5/SIP1, PCK1, MGL;* Fig. 3A, cluster 3) and *GolS2, BAM1/BMY7, AVP1/AVP3, HKL3* specifically in roots; Fig. 3A, cluster 5), ROS scavenging (e.g. *GSTU19/GST8;* Fig. 3A, cluster 3) and *GPX3*; Fig. 3A, cluster 5), dehydrins (e.g. *LTI29/ERD10/LTI45;* Fig. 3A, cluster 3) and *LEA7, LEA4-1* specifically in roots; Fig. 3A, cluster 5), ABA biosynthesis and signalling (e.g. *NCED3/SIS7/STO1, NCED5, PP2CA/AHG3, CIPK6/SNRK3.14/SIP3, AFP2;* Fig. 3A, cluster 3). Additionally, genes that restrict proline catabolism (e.g. *ERD5/PDH1, PDH2;* Fig. 4A, cluster 10), ROS production (e.g. *RBOHF, RBOHD;*Fig. 4A, cluster 9 and *GSTF10/ERD13, GSTF6/GST1/ERD11/GSTF3, GST30/ERD9/GSTU17;* Fig. 4A, cluster 10) and auxin and cytokinin signalling (e.g. *UGT74E2;* Fig. 4A, cluster 8 and *UGT76C2, WOL/CRE1/AHK4;* Fig. 4A, cluster 10) displayed conserved transcriptional responses. While this confirmed overlap in the stress-response between the species, especially for osmotic and oxidative tolerance, notable differences were observed in the transcriptional response of TFs.

**Table 2.** Drought responsive genes of *Arabidopsis thaliana* identified as *Picea abies* homologs and orthologs.

| Plant species | Genome | Needles | Roots | Common |
| --- | --- | --- | --- | --- |
|  |  | Transcriptome/<br>DE | Transcriptome/<br>DE |  |
| *Arabidopsis*<br>(373 genes) | 335<br>(89.8 %) | 325/39<br>(87.1/10.5 %) | 328/117<br>(87.9/31.4 %) | 322/33<br>(86.3/8.9 %) |
| Norway spruce | 979 | 782/43 | 826/164 | 758/24 |

### Identification of TFs involved in regulating drought stress in Norway spruce

Among the DE genes in needles and roots were 74 and 240 TFs, respectively (Fig. 5). These TFs included members of 20 and 33 TF families, respectively, from the 54 TF families annotated in the Norway spruce genome (Fig. S4 and Supplementary file 4). Among all TF families there was significant enrichment (p < 0.05) of ERF, C2H2, MYB-related and NAC TF families in needles and ERF, MYB and SAP in roots (Fig. 5 & S4). Four TFs in needles and 26 in roots had a known drought function in Arabidopsis (Supplementary file 3 & 4), three of which were DE in both tissues. TFs of the ABA-dependent pathway, which are typically observed to be drought-responsive in Arabidopsis (the AREB/ABFs), were, however, absent. Instead, ABA-independent signalling via *DREB2B* and *ERF53* was more pronounced in Norway spruce, particularly in roots (see below), indicating that different regulatory pathways are active in Norway spruce seedlings in response to drought. In needles, TF ZF2 was up-regulated in cluster 2 (Fig. 3A). In cluster 3, NAC TFs (NAC3/NAC055, RD26) and ERFs (DREB19, ERF7) were commonly up-regulated in both tissues (Fig. 3A). In roots, the earliest transcriptional response was up-regulation of MYB2 in cluster 4 after five days, the expression of which remained high under severe water stress (Fig. 3A). Severe drought in roots additionally activated NAC019, TFs within the ERF family (DREB2B, ERF53, RAP2.1, RAP2.4, RAP2.6L), a WRKY TF (WRKY33) and MYBs (MYB96, MYBR1) (Fig. 3A, cluster 5). ARR-B TFs (RR10, RR1) and ZFHD1/ZFHD11/HB29 were also up-regulated after severe drought in roots. Down-regulation of TFs in roots after severe drought occurred for ERF family members (CBF1/DREB1B, CBF3/DREB1A, CBF4/DREB1D, HRD, DDF1, RAP2.1, SHN1/WIN1; Fig. 4A, cluster 10). Also, MYB TFs (MYB61, MYBR1), a BHLH TF (BHLH100) and a HD-ZIP TF (HB6) were down-regulated. After re-irrigation, increased expression of TFs, such as of MYB15, RAP2.6 or RAP2.6L and NAC3/NAC055 and RD26 was found in roots (Fig. 3A, cluster 6 and 7). Among the drought-responsive TFs were a number of Norway spruce-specific singletons (Supplementary file 4). As such, these observed transcriptional responses to drought for the Norway spruce orthologs and homologs of the considered set of well-characterised Arabidopsis genes showed evidence of both conservation and substantial divergence of transcriptional response between these two species.

**Figure 5.**
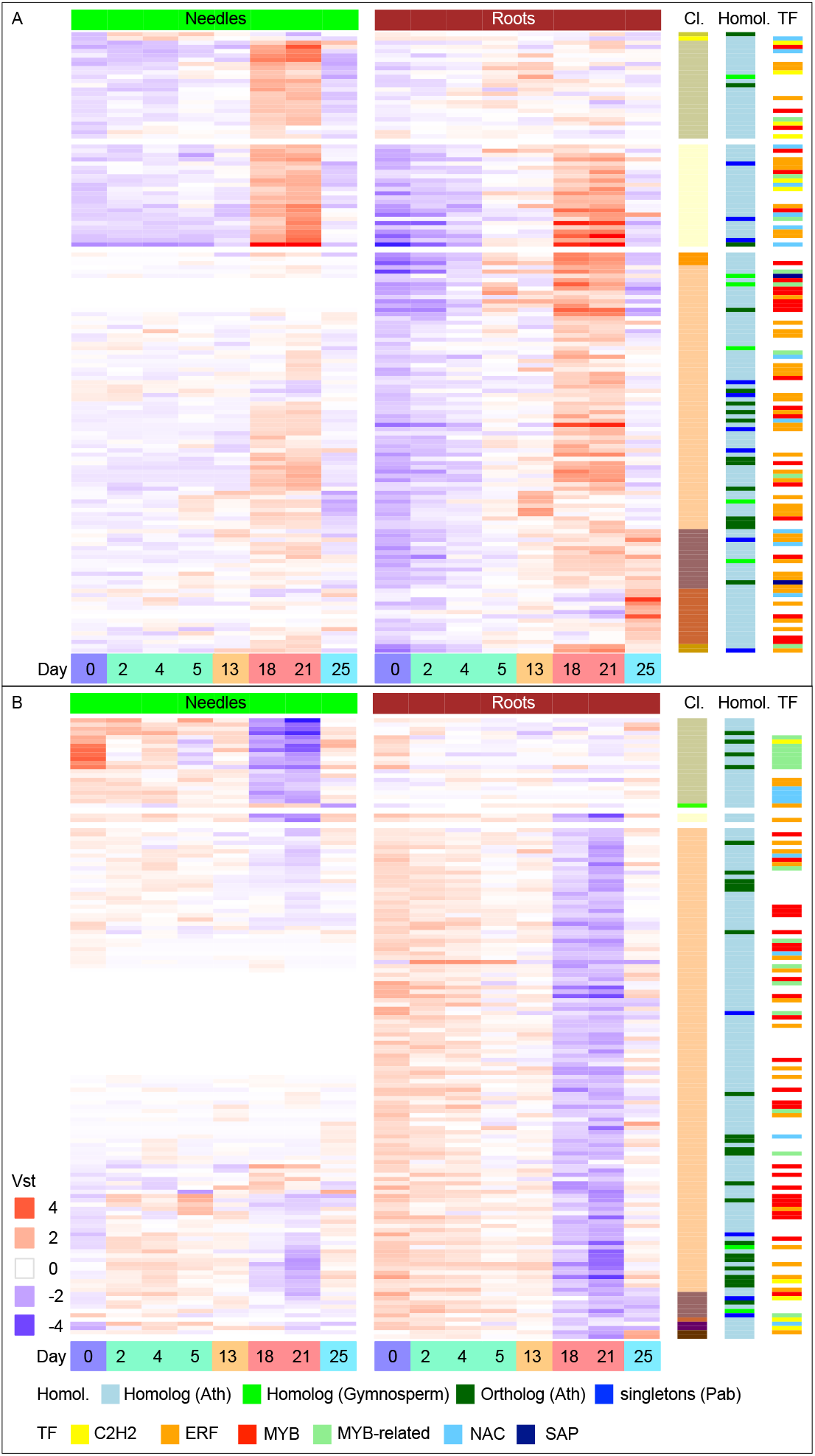
Heatmap diagram of transcription factors differentially expressed in Norway spruce in response to drought. Displayed are the differentially expressed transcription factors over the eight sampling time points (day 0, 2, 4, 5, 13, 18, 21 and 25) and separated by tissue, needles (green) and roots (brown). Time points were coloured corresponding to Figure 1 in blue at day 0, green at day 2,4 and 5, orange at day 13, red at day 18 and 21 and turquoise at day 25. A) Up-regulated transcription factors were clustered (Cl.) by expression and sorted by the same colour coding as in Fig. 3A. B) Down-regulated transcription factors were clustered by expression and sorted by the same colour coding as in Fig. 4A. Displayed are the vst values scaled by row means. In addition, information on sequence homology (Homol.) is visualised on the right side, with genes orthologous to Arabidopsis coloured dark green, homologous to Arabidopsis in light blue, homologous to other gymnosperms in light green and Norway spruce specific singletons in dark blue. Furthermore, transcription factor (TF) families significantly enriched: ERFs, MYB-related, C2H2s and NACs in needle samples and ERFs, MYBs and SAPs in roots are colour coded.

It should be noted that exact ortholog inference is currently challenging for Norway spruce due to the fragmented nature of the genome assembly (Nystedt et al., 2013). As such, caution is needed in interpreting inference of functional ortholog conservation. This likely underlies many of the cases where multiple copies of the same functional gene were identified, as in the case of ABCG40 (Supplementary file 3), for which four copies were identified as expressed in Norway spruce roots. In some such cases there were differences in which gene copy was DE between the tissues. For example, there were two orthologs of the potassium channel protein KT1, which were DE between the two tissues. While this potentially represents two genuine copies that have undergone sub-functionalisation, such observations require detailed follow-up studies or reconsidering upon availability of an improved genome assembly.

## Discussion

Drought stress typically induces characteristic, short-term physiological and molecular responses that enable plants to survive periods of limited or fluctuating water availability. Drought stress responses are initiated by limitation of soil water uptake by roots, with root-level responses to this initial limitation being important determinants of drought tolerance and survival. Despite the importance of root responses, the majority of previous studies have focused on the response of aboveground tissues. In this study, drought-induced transcriptional changes of both roots and needles of Norway spruce seedlings were assayed by subjecting seedlings to a progressively increasing soil water deficit, followed by a recovery phase after re-irrigation. The induced transcriptional responses were placed within a framework of physiological responses observed in needles. To explore the extent of conservation in the transcriptional drought response, the transcriptional response of Norway spruce was compared to previous results in Arabidopsis, as the most comprehensively studied model system for drought and other abiotic stress responses.

g_s_ and A_max_ were at their observed minimums at day 18 and midday shoot water potential was as low as −2.1 MPa at day 21, before which there was little observed physiological response to water stress in the needle samples (Fig. 1D). In a similar study of red spruce seedlings (*Picea rubens* Sarg.) photosynthesis continued until water potential decreased to −3.0 MPa and was rapidly recovered to 70% of well-watered plants after re-irrigation (Seiler and Cazell, 1990). In the current experiment re-irrigation after 21 days resulted in 80% recovery of the photosynthetic rates (Fig. 1C). Nevertheless, the treatment was regarded as severe, with death occurring in seedlings that experienced an additional two days of severe drought. In agreement with this observation, it was reported earlier that Norway spruce seedlings die after crossing a critical point of dehydration (Ditmarova et al., 2010). These findings highlight the severe negative effects that could result from changing patterns of soil water availability as a result of continued climate change and the associated increasing occurrence of severe weather events, including extended periods of limited precipitation. As such, there is a clear need to further advance our understanding of the drought responses of Norway spruce and related species to both predict responses and adaptive potential to such future conditions and to inform breeding and development programs to generate new varieties that can survive and maximise productivity under those conditions. As an example from the current study, for breeding of more drought tolerant Norway spruce seedlings knowledge of root-specific and early-response genes, such as the *MYB2* TF, represent potential new targets for downstream functional studies and selection programs.

The limited physiological response of needles to severe drought was reflected in the transcriptome, with substantially fewer DE genes in needles than in roots (Fig. 2C). In contrast to the needle response, changes in the transcriptome of roots to an increasing soil water deficit were observed starting at 30% FC, with the response being particularly prominent in response to severe drought (day 18; Fig. S1B). Correspondingly, a greater number of GO biological processes were represented and significantly enriched among DE genes in roots. Nevertheless, there was a common enrichment in both tissues of up-regulated genes for “response to stress”. The extensive down-regulation in roots was enriched for a number of categories associated with growth (e.g. “anatomical structure development”, “growth”, “cell wall organisation or biogenesis”), metabolism (e.g. “carbohydrate metabolic process”) and “transport” related processes, indicating that reduced root development and growth is an important component of the drought response mechanism in the roots of Norway spruce seedlings. In contrast, the majority of DE genes in needles were up-regulated, with representation of genes in categories including “transport”, “small molecule metabolic process”, “cellular amino acid metabolic process”, “signal transduction”, “reproduction”, “biosynthetic process” and “anatomical structure development” during severe drought, indicating that metabolism similar to pre-stress is maintained in needles. In agreement, there was no enrichment for terms associated with photosynthesis, suggesting that needles, in contrast to roots, did not adjust their primary physiological mechanisms via modulation of the transcriptome and corroborating that both physiological and molecular responses to drought stress are limited in needles before severe damage and needle loss occurs.

Comparison of transcriptional regulation between species is important to determine the evolution of stress responses and to ascertain the transferability of biological knowledge across species. Here, indications of both conservation and substantial divergence in transcriptional responses were observed between Arabidopsis and Norway spruce. The conserved component of the Norway spruce response included up-regulation of solute accumulation, ROS scavenging, ABA biosynthesis and signalling. In contrast, proline catabolism, ROS production and auxin and cytokinin signalling responses were limited. A large proportion of homologs or orthologs of the characterised Arabidopsis drought responsive genes considered were, however, not DE in response to drought, suggesting that there are extensive, non-conserved, components of the transcriptional drought response between the two species. Of note, ABA-dependent regulation was less pronounced in Norway spruce than in Arabidopsis. Whether this reflects a general divergence of the drought response mechanisms between lineages or is a specific component of contrasting (isohydric versus anisohydric) drought response strategies requires further studies. In addition to these conserved and diverged components of homologous genes present in both genomes, there were additionally a number of Norway spruce-specific genes that were under active regulation in response to drought. These genes are uncharacterised and lack functional annotation on the basis of homology, making them interesting targets for future characterisation studies.

Differences and parallels between species were, among others, identified in the transcriptional response of genes within the ABA signalling pathway. Sensing and signalling by plant hormones plays an important role in the early response to drought stress and production of ABA is a well-recognised component of the typical response to water deficit (Schachtman and Goodger, 2008). In the current study, up-regulation was observed in needles and roots for the ABA-synthesis genes *NCED3* and *NCED5*, a *PP2CA* gene and of the serine/threonine kinase *SnRK2*, involved in ABA signalling. However, members of the central TFs of the ABA-dependent transcriptional activation under drought stress, the AREB/ABFs (Singh and Laxmi, 2015), were not DE. While expression of homologs to *AREB1/ABF2, AREB2/ABF4* and *ABF3* increased, these changes were not significant. In contrast, TFs of the ABA-independent DREB1/CBF and DREB2 regulons, DREB1A/CBF3, DREB1B/CBF1, DREB1C/CBF2 and DREB2B, were significantly DE in roots. *DREB1/CBF* type genes have mainly been reported to be responsive to low temperatures in Arabidopsis (Nakashima et al., 2009) but were also up-regulated during water deficit in maize shoots and, to an even greater extent, in roots (Liu et al., 2013), with similar expression changes reported in poplar and loblolly pine roots (Cohen et al., 2010; Lorenz et al., 2011). In the current study, expression of *DREB1/CBF* genes was strongly down-regulated in roots and only *DREB2B*, which has a well-defined function in drought stress (Nakashima et al., 2009), was up-regulated in roots. Similarly, expression of *ERF53* increased under drought stress, which has been proposed to function in a similar manner to *DREB2A* via regulation of drought stress-responsive genes (Cheng et al., 2012). Together, these findings indicate that Norway spruce seedlings use a different set of regulators to control the expression of drought responsive genes compared to the typical response of Arabidopsis, with the ABA regulated signalling pathway playing a less important role than in many angiosperm species.

While many genes differed in their response to drought between Arabidopsis and the current Norway spruce seedlings, a number of genes with known functional roles in drought tolerance were DE and showed a transcriptional response in needles and roots, suggesting conserved function. The observed drought response involved expression-conserved up-regulation of solute synthesis, such as of malate (PCK1) (Penfield et al., 2012), raffinose (RS5/SIP1, GolS1) (Taji et al., 2002; Nishizawa et al., 2008; Egert et al., 2013; Rasheed et al., 2016) or amino acids (MGL) (Joshi and Jander, 2009) and protection of cellular components via expression of dehydrins (LTI29/ERD10/LTI45) (Kasuga et al., 2004) and ROS scavengers (GSTU19/GST8 and GST30/ERD9/GSTU17) (Chen et al., 2012; Xu et al., 2016) in both tissues. Furthermore, other regulatory genes had a conserved function in Norway spruce. For example NAC and MYB TFs, such as ANAC072/RD26, which were here up-regulated in needles and roots, have been shown to improve drought stress tolerance in Arabidopsis when overexpressed (Nakashima et al., 2012). The *ANAC019* and *ANAC072/RD26* genes, here with root-specific expression, have been classified as stress-responsive NAC (SNAC) TFs (Tran et al., 2004). Expression changes of many MYB TFs have been reported during drought stress responses in plants (Baldoni et al., 2015), although their functional role has not been elucidated. *MYB2* is known to mediate response to abiotic stress and regulate gene expression of ABA-inducible genes (Abe et al., 2003) and to be expressed in both shoots and roots of Arabidopsis and rice (Yang et al., 2012; Baldoni et al., 2015). This gene was DE early on in roots in the current study, suggesting functional conservation. Severe drought stress induced *MYB96* in roots, which may function to arrest lateral root growth in a similar manner to the described role in Arabidopsis (Seo et al., 2009). This response is suggested to enable maximal resource allocation to primary root growth to facilitate rapid recovery after cessation of water shortage (Comas et al., 2013). During recovery, *MYB15* was here expressed in roots and, in Arabidopsis, this gene has been reported as a negative regulator of *DREB1/CBF* genes during cold stress (Agarwal et al., 2006). Potentially this gene may therefore have a similar function in Norway spruce prior to and during recovery.

## Conclusion

This study provides a comprehensive characterisation of the transcriptional drought response mechanisms in Norway spruce roots and needles, serving as a guide for future studies, including breeding and selection of targets, further deconvolution of the regulatory control of drought stress response and the adaptive capacity of Norway spruce seedlings under future climate conditions. The transcriptional response to drought of Norway spruce seedling roots and needles differed substantially. There were extensive changes in gene regulation in roots, with down-regulated modulation of genes related to growth and metabolism. In contrast, little transcriptional response was observed in needles, reflecting the limited physiological changes in water potential and stomatal conductance shown until severe drought was experienced. This shows that Norway spruce has particularly contrasting transcriptional response mechanisms in above- and belowground tissues and that active modulation of the root transcriptome is an important component of the drought response. Comparison to a curated set of well-characterised drought responsive genes in Arabidopsis revealed examples of Norway spruce genes with expression-based evidence of a conserved role in drought response. More generally, however, the observed transcriptional response of a larger proportion of Norway spruce orthologs suggested divergence of function, particularly so for genes within the ABA-dependent drought response pathway.

## Material and Methods

### Experimental design

Three year old Norway spruce (*Picea abies L*. (Karst.) seedlings of the seed provenance Lilla Istad (56° 30’ N) were grown in a growth room at 18 h of light (120 μmol m^-2^ s^-1^ at plant height) and 20 °C in 3-l pots filled with peat. For a controlled drought treatment the FC of the soil was determined as the difference in weight of wet soil and dry weight, after oven drying at 70 °C. Seedlings of the control treatment were kept well-watered and monitored gravimetrically to maintain soil moisture at 80% FC. For a mild drought treatment, water was withheld from the seedlings until soil moisture reduced to 30% FC, five days after start of the experiment. Seedlings were kept in this stage of mild drought stress for seven days by adding the daily evapotranspirational water loss. This was a non-lethal drought stress and above a point where leaf death would occur. A severe drought stress was then imposed by completely withholding water until symptoms of severe dysfunction, as measured by photosynthetic assimilation rates, were observed. At this moment the severe drought stress was extended for three days, bringing plants closer to catastrophic hydraulic failure, before rehydration was started. After four days of re-watering soil moisture had returned to the initial well-watered conditions at 80% FC. Needles and fine and small lateral roots (<4 mm) of three to four seedlings were sampled at control (day 0), mild (2, 4, 5 and 13 days) and severe (18 and 21 days) drought and after re-irrigation (25 days) for RNA-Seq described below.

### Gas exchange and water potential measurements

Gas exchange measurements were performed with a portable infrared gas analyser (model LI-6400XT; Licor, Lincoln, NE, USA) at days 0, 2, 4, 5, 13, 16, 17, 18 and 25 of the drought treatment. Leaf photosynthesis at light saturation (A_max_) and g_s_ were measured between 10:00 and 15:00 on three control and three drought treated plants not used for RNA-Seq in a randomised sequence. Photosynthetic photon flux density inside the cuvette was maintained at 1500 μmol m^-2^ s^-1^ and CO_2_ concentration at 400 μmol mol^-1^. Further, leaf-to-air vapour pressure deficit was between 1-1.5 kPa, leaf temperature was close to 22 °C and humidity in the cuvette was maintained above 60%. Photosynthetic leaf area of the shoots used for gas exchange measurements was determined using ImageJ (Schneider et al., 2012; Rueden et al., 2017). Water potential (Ψ_s_hoot) was measured at midday on three lateral shoots from each of three-five plants on day 0, 2, 4, 5, 13, 18, 21 and 25 of the treatment. The cut shoots were sealed in a plastic bag and stored in an insulated container until measurement with a Scholander-type pressure chamber (SKPM 1400, Skye instruments Ltd., Powys, UK). Visualisation and statistical analysis was performed in R 3.4.3 (R Development Core Team, 2018). Values for A_max_ and g_s_ were set relative to day 0. Differences in A_max_ or gs over time in the treated plants were tested with one-way repeated measure analysis of variance (ANOVA) using the package nlme (Pinheiro et al., 2018). Posthoc tests were performed using the multcomp package (Hothorn et al., 2008) with Tukey’s honestly significant difference (HSD) test. Differences in water potential were assessed with a one-way ANOVA and time as the factor. A per-time comparison *t*-test was then applied to test significant differences between measurements of control plants and water-stressed plants. Significant differences were assigned at P < 0.05.

### RNA extraction and sequencing

The same plants used for water potential measurements were sampled for analysis of the drought transcriptome. At day 0, 2, 4, 5, 13, 18, 21 and 25 of the treatment needles and roots of three to four seedlings were collected on dry ice and stored at −80 °C until further processing. They were ground manually in liquid nitrogen and total RNA was prepared after Chang et al. (1993) with the following modifications: Addition of warm extraction buffer, including polyvinylpyrrolidinone (PVP) 40, and vortex mixing of the ground sample material was followed by an incubation step for 5 min at 65 °C. Precipitation with ¼ volume 10 M LiCl took place at −20°C for 2 h and RNA was then isolated by centrifugation at 14.000 rpm for 20 min and 4 °C. The RNA was further purified using the RNeasy mini kit (QIAGEN, Hilden, Germany), following the manufacturer’s instructions. A DNase digestion with the RNase-free DNase set (QIAGEN) was performed. Finally the RNA was eluted in 40 μl of RNase-free water for 2 min at room temperature. Integrity of total RNA was assessed with the Agilent RNA 6000 Nano kit (Agilent Technologies, Waldbronn, Germany) on a Bioanalyzer 2100 (Agilent Technologies) and purity measured with a NanoDrop 2000 spectrophotometer (Nanodrop Technologies, Wilmington, DE, USA). High quality total RNA with a RIN ≥ 7.5, OD 260/280 ratio of ≥ 2.0 and concentrations ≥ 50 ng/μl was sequenced by SciLifeLab (Stockholm, Sweden) for paired-end (2 x 125 bps) sequencing on a HiSeq 2000 platform using standard Illumina protocols. The sequencing library preparation included an enrichment for poly-adenylated mRNAs and all samples yielded > 12.4 million read pairs.

### Sequencing data analysis

Raw reads were pre-processed following a guideline for RNA-Seq data analysis implemented in the bioinformatics platform at Umeå Plant Science Centre (https://bioinformatics.upsc.se/publications) (Delhomme et al., 2014). Briefly, after an initial quality assessment of the raw data with FastQC (version 0.11.2; https://www.bioinformatics.babraham.ac.uk/projects/fastqc/) ribosomal RNA was removed with SortMeRNA (version 2.0) (Kopylova et al., 2012). Sequencing adapters and low quality regions were cut using Trimmomatic (version 0.36) with the setting ILLUMINACLIP:2:30:10 (Bolger et al., 2014). Reads were then aligned to the Norway spruce reference genome (*Picea abies* v1.0) using STAR (version 2.4.0f1) set to –outSAMmapqUnique 254 –quantMode TranscriptomeSAM – outFilterMultimapN-max 100 –chimSegmentMin 1 (Dobin et al., 2013) and a count table generated with HTSeq (version 0.6.1) and settings –m intersection-nonempty –s yes –t exon –i Parent (Anders et al., 2015).

Count tables of needle and root data were filtered separately for genes with more than one sequencing read in at least two biological replicates, leaving 43639 needle and 47880 root genes in the tissue specific drought stress transcriptomes. In a next step the variance stabilisation transformation (vst) function of DESeq2 (version 1.16.1) (Love et al., 2014) was used to normalise expression of the data sets in R. Visualisation by principal component analysis (PCA) was used to provide a visual overview of the treatment effect on the transcriptome and enabled identification of outlier samples for removal from further analysis (one of the needle samples at day five).

On the basis of the PCA plot, samples were grouped into mild (2,4 and 5 days after start of the experiment), severe (day 18 and 21 of the treatment) stress and rehydrated samples (day 25) for calling DE genes and pairwise comparison between the groups and well-watered samples at day 0 for each tissue separately were performed using DESeq2. Results were filtered for a Log2 fold change > 2 and an adjusted *p* value < 0.01.

Gene Ontology (GO) Slim annotation and enrichment analysis was performed on clusters of differentially expressed genes identified by ComplexHeatmap analysis (Gu et al., 2016) in R after calculating the mean of the expression for each sampling point on vst values of the count tables. Default method for clustering was “hclust” and “euclidean” distance and vst values were scaled by row means. 13 Clusters separated up- and down-regulated genes by tissue and time point of expression. Gene Ontology annotations were obtained from the Conifer Genome Integrative Explorer (ConGenIE; http://congenie.org) (Sundell et al., 2015). GOSlim annotations were built for Norway spruce by the Map2Slim function of Owltools (https://github.mm/owlmllab/owltools/wiki/Map2Slim),using the plant slim subset provided by the Gene Ontology Consortium (Ashburner et al., 2000; Beike et al., 2015). For enrichment analysis the program GeneMerge (Castillo-Davis and Hartl, 2003) was used and the background of expressed genes adjusted, with a vst value cut-off of > 4 (transcriptome size needles 42821, roots 47880 genes), and enrichment assigned if P < 0.05.

Sequence homology BLASTp searches of the differentially expressed *Picea abies* gene models were performed against *Arabidopsis thaliana*, other gymnosperms (*Picea glauca*, *Picea sitchensis*, *Pinus pinaster*, *Pinus sylvestris*, *Pinus taeda*, *Pseudotsuga menziesii*, *Gnetum montanum*, *Taxus baccata*, *Cycas micholitzii* and *Gingko biloba*) and further angiosperm species (*Populus trichocarpa, Oryza sativa ssp. japonica* and *Amborella trichopoda*) inclusive *Physcomitrella patens*. Orthologs were detected through OrthoMCL and homologs using highest sequence homology results as provided by Gymno PLAZA 1.0 (Proost et al., 2015). Previously recognised genes functioning in drought response in *Arabidopsis thaliana* from the DroughtDB (Alter et al., 2015) and from drought related GO terms: “response to water deprivation” (GO:0009414), “cellular response to water deprivation” (GO:0042631), “regulation of response to water deprivation” (GO:2000070) and all subcategories linked with these were used for a collective search of conserved function within the differentially expressed transcripts in samples of needles and roots of Norway spruce.

TFs and their corresponding families were obtained from PlantTFDB 3.0 (Jin et al., 2014). Hypergeometric tests were performed using the “phyper” function in the base R package stats and enrichment for TF families in the background of all TF in the transcriptomes and of singletons (if no sequence homology of any kind existed) in the set of DEG in comparison to the transcriptome sizes calculated and significance assigned if P < 0.05.

Sequencing data has been deposited at the ENA (European Nucleotide Archive, https://www.ebi.ac.uk/ena) under the accession ID PRJEB26933.

## Availability of data and materials

The datasets generated and/or analysed during the current study were deposited to the European Nucleotide Archive (ENA, www.ebi.ac.uk/ena) and are available under the accession number PRJEB26933.

## Acknowledgements

The authors wish to thank the UPSC bioinformatics facility (https://bioinfomatics.upsc.se) for technical support during the RNA-Seq data pre-processing and analyses, the greenhouse staff at Umeå Plant Science Centre for horticultural services, as well as Catherine Campbell for support with the gas exchange measurements.

## Supplemental Data

**Supplemental Figure 1 Principle Component Analysis plot of needle and root gene expression data during progressing drought.**

Expression data of A) needles and B) roots were normalised using variance stabilising transformation before ordination analysis. Three to four seedlings were sampled at control, mild and severe drought conditions and after rehydration and used for RNA sequencing. Displayed are the first two components of the PCA with samples coloured by soil water percentage field capacity.

**Supplemental Figure 2 Venn-diagrams of differentially expressed genes in response to drought treatments.**

Differentially expressed genes in needles (left) and roots (right) separated in up- (top) and down-regulated (bottom) genes. Numbers represented the genes differentially expressed in seedling of the mild (yellow) and severe (dark green) drought against the control or after re-irrigation (gray).

**Supplementary Figure 3. Heatmap diagram of Norway spruce genes with a conserved drought function.**

Spruce genes with a conserved function in drought in Arabidopsis were clustered by expression and displayed by sampling time points. Time points were coloured corresponding to Figure 1 in blue at day 0, green at day 2,4 and 5, orange at day 13, red at day 18 and 21 and turquoise at day 25. The upper heatmap displays all genes found in the needle transcriptome, the lower heatmap shows included genes of the root transcriptome after removal of genes with low read counts (at least one read in two biological replicates). Differentially expressed genes are additionally indicated with black lines on the right side (DE). Displayed are the vst values scaled by row means.

**Supplemental Figure 4. Bar graph of differentially expressed transcription factors in needles and roots of Norway spruce in response to drought.**

Differentially expressed transcription factors are distributed by transcription factor families and scaled by the total number of differentially expressed transcription factors in a family. A) Transcription factors were found in needles (green), roots (brown) or were in common (violet/rose) for the tissues. They were either up- (light colour) or down-regulated (dark colour) or expression could change during progression of the treatment (“other”; gray). B) Furthermore, the transcription factors were either orthologous (dark green) or homologous (light blue) to Arabidopsis transcription factors or were homologous to other gymnosperms (light green) or Norway spruce specific singletons (dark blue).

**Supplementary file 1. Summary of differentially expressed genes during drought in Norway spruce.**

Genes differentially expressed in needles, roots or both tissues are listed and sorted by cluster membership as assigned on the basis of expression trends. Additional information is available on Gene Ontology Slim categories, Transcription Factor families and homology to other species.

**Supplementary file 2. Gene Ontology Biological processes categories of the drought response in Norway spruce.**

Listed were the Gene Ontology Slim categories of biological processes induced (Sheet 1) or repressed (Sheet 2) in drought stressed Norway spruce samples and the number of differentially expressed genes present within each cluster.

**Supplementary file 3. Differentially expressed genes with a conserved function in drought.**

Genes with a known function in Arabidopsis drought response and their corresponding orthologs or homologs in Norway spruce are listed. Sheet 1 details genes differentially expressed in Norway spruce, while Sheet 2 contains the full list of selected Arabidopsis genes and whether their corresponding Norway spruce orthologous/homolgous genes were differentially expressed.

**Supplementary file 4 Transcription factors regulated during drought in Norway spruce**

Number of genes categorized as differentially expressed transcription factors in Norway spruce. For each transcription factor family the numbers of differentially expressed transcription factors in a tissue, up- or down-regulated, is given and homology information provided.

Author contributions
JCH, VH and NRS planned and designed the study. JCH performed the experiments and analysed the data together with AV, VH and NRS. JCH wrote the initial draft of the manuscript. JCH, AV, VH and NRS edited the manuscript into its final form. All authors read and approved the final manuscript.

## Notes

**Funding:** This work was supported by funding from HolmenSkog AB and Berzelii Centre for Forest Biotechnology to VH, and from the Swedish University of Agricultural Science’s Trees and Crops for the Future (TC4F) program to VH and NRS.

